# Accurate structure prediction of immune proteins using parameter-efficient transfer learning

**DOI:** 10.1101/2024.11.13.621715

**Authors:** Tian Zhu, Milong Ren, Zaikai He, Siyuan Tao, Ming Li, Dongbo Bu, Haicang Zhang

## Abstract

Accurate prediction of immune protein structures is crucial for understanding the immune system and advancing immunotherapy development. While deep learning methods have significantly advanced protein structure prediction by extracting evolutionary constraints from homologous sequences of a target protein, they struggle with immune proteins due to the limited number of known structures and the lack of homologous sequences in hypervariable regions. To address this challenge, we propose ImmuneFold, a transfer learning approach that fine-tunes ESMFold specifically for immune proteins. We leverage low-rank adaption (LoRA), a parameter-efficient fine-tuning technique that requires considerably less memory and substantially fewer parameters. Evaluations on various immune proteins, including T-cell receptors, antibodies, and nanobodies, demonstrate that ImmuneFold outperforms existing methods in prediction accuracy. Furthermore, we apply ImmuneFold to develop a zero-shot protocol for TCR-epitope binding prediction. Unlike previous supervised methods suffering from severe overfitting due to limited experimental binding data, our approach first predicts TCR-epitope structure using ImmuneFold and then directly estimates the binding affinity by calculating Rosseta energy. Evaluations on experimental binding datasets suggest that our method is robust and accurate in predicting TCR-epitope binding. In summary, ImmuneFold demonstrates accurate predictions of immune protein structures and TCR-epitope binding, highlighting its potential to advance the development of immunotherapies.

## 1 Main

T-cell receptors (TCRs) and antibodies are critical proteins in the adaptive immune system, responsible for recognizing and neutralizing specific antigens. TCRs specialize in targeting neoantigens and foreign peptides presented by major histocompatibility complexes (MHCs), while antibodies can bind to the surface of a wide range of antigens. Both have been extensively utilized in the development of personalized immunotherapies and vaccines [1–3]. For example, over a hundred antibody drugs have been approved, and the first TCR-based cancer therapy was recently authorized [4]. Additionally, nanobodies—single-domain antibodies found in camelids —have smaller molecular sizes and offer unique advantages in drug development [5].

All these three types of immune proteins belong to the immunoglobulin (Ig) superfamily. Structurally, their framework regions are conserved and maintain the overall architecture, while the complementarity-determining regions (CDRs) are highly variable in both structure and sequence, primarily determining antigen-specific binding. Upon binding to specific antigens, antibodies, and TCRs form antibody-antigen complexes and TCR-pMHC complexes, respectively. Understanding the structural basis and antigen-specific binding mechanisms of these complexes can enhance structure-based drug design with precise binding specificity and affinity. However, experimental methods like X-ray crystallography and cryo-electron microscopy (cryo-EM) are costly and require pure, stable protein samples. Therefore, there is increasing interest in computational modeling of immune protein structures, especially focusing on the CDRs.

While recent deep learning methods, such as AlphaFold [6] and RoseTTAFold [7] have significantly advanced protein structure prediction, accurate prediction of immune protein structures remains challenging. These methods rely on modeling the evolutionary constraints in the multiple sequence alignments (MSAs) of homologous sequences, thus their predictive accuracy is contingent upon the quality of MSAs. However, the CDRs, particularly CDR3 in the antibody heavy chain or TCR *β* chain, exhibit high variability and lack sufficient homologous sequences.

To address the limitations of MSA-based methods, language model-based approaches specifically designed for antibody structure prediction have been proposed [8, 9]. Rather than relying on homologous sequences, these methods extract the evolutionary constraints from large-scale language models trained on extensive antibody sequences datasets [10, 11]. They have demonstrated comparable or superior performance while significantly improving efficiency, with structures predicted in seconds. However, the prediction accuracy of these methods is constrained by the scarcity of structure data as only thousands of antibody structures are currently available in the Protein Data Bank (PDB). The challenges are even more pronounced for TCRs. Although millions of CDR sequences are available, the number of full-length TCR variable domain sequences is limited, making it difficult to train effective language models for structure prediction. Furthermore, only a few hundred TCR structures are available for training, which is orders of magnitude fewer than for antibodies.

We propose ImmuneFold, a transfer learning approach that fine-tunes ESMFold [12] on immune protein structures. This method enables the model to leverage knowledge learned from massive protein structures beyond immune proteins, thereby alleviating the issue of limited training data. Specifically, we employ the low-rank adaptation (LoRA) technique for fine-tuning, which updates only a subset of parameters rather than the entire model. Compared with full-parameter fine-tuning, LoRA requires significantly less memory and training time while maintaining performance [13–15], making it more accessible for academic research groups with limited computational resources. Originally, LoRA is proposed to fine-tune the Transformer architecture in large-scale language models like Llama2 [13]. It has since been extended to the image generation task [16] and more recently applied to protein language models for various protein-related tasks [15, 17–19]. In this work, we extend LoRA to the Evoformer-like architecture, a crucial yet computationally expensive component of recent deep learning models for protein structure prediction.

To demonstrate the practical utility of ImmuneFold, we apply it to develop a protocol for predicting TCR-epitope binding. Accurate prediction of TCR-epitope interactions is essential for selecting peptides that trigger immune responses and for engineering TCRs capable of efficiently recognizing specific neoantigens in immunotherapy [20]. However, existing supervised methods [21–23] suffer from severe overfitting due to the scarcity and biased distribution of training binding data. They often fail for TCRs absent from training data, limiting their utility in the practical immunotherapy application. To address this challenge, we integrate the ImmuneFold predictions with Rosseta energy [24] to directly evaluate TCR-epitope binding affinity in a zero-short manner, thereby avoiding the need for training binding data.

## 2 Results

### 2.1 Low-rank adaption of ESMFold for immune protein structure prediction

Within the transfer learning framework capable of alleviating the issue of limited training data, ImmuneFold leverages the Low-Rank Adaptation (LoRA) technique to fine-tune ESMFold for immune protein structure prediction (Figure 1).

**Fig. 1:**
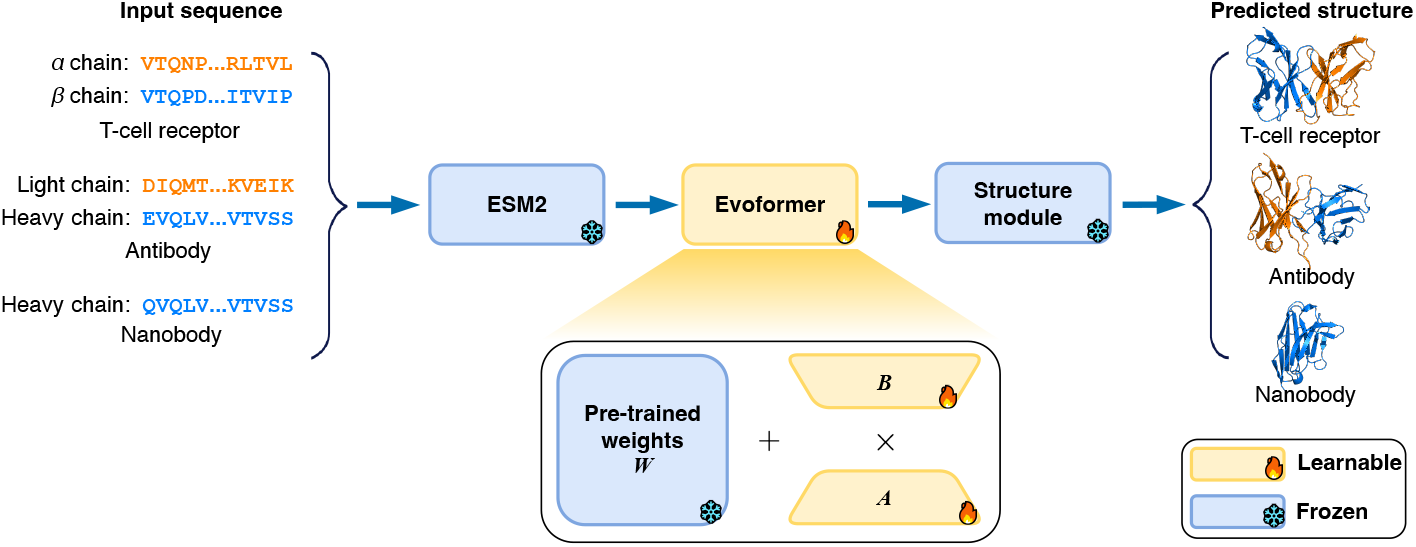
Overview of ImmuneFold. ImmuneFold employs Low-Rank Adaptation (LoRA), a parameter-efficient fine-tuning technique, to fine-tune only the Evoformer module of ESMFold on immune protein structures. Two separate models are fine-tuned: one for TCR structure prediction and the other for antibody and nanobody structure prediction.

Our approach offers two main advantages. First, we initialize the model weights from ESMFold, enabling the use of learned presentations from massive protein structures beyond immune proteins. Second, LoRA updates only two low-rank matrices instead of the entire weight matrix, significantly reducing the number of trainable parameters [13, 25]. Compared to full-parameter fine-tuning, which requires extensive computational resources, LoRA is more accessible for academic research groups without sacrificing prediction performance [16, 26].

We apply LoRA solely to the `Main Folding Trunk` module of ESMFold, an Evoformer-like architecture that learns single and pair representations and serves as a key component for protein structure prediction [6]. We keep the other two modules frozen: the ESM2 language model for sequence embedding and the `Structure Module` that maps the learned representations to 3D coordinates.

To accommodate various real-world application scenarios, ImmuneFold supports structure prediction for both unbound immune proteins and their bound states when antigen context is provided. We trained two separate models for TCRs and antibodies, respectively.

### 2.2 Accurate TCR structure prediction with ImmuneFold

We derive the training data for TCR structure prediction from the STCRDab database [27] and evaluate performance using independent testing sets with no overlap with the training data. We compare our approach with TCR-specific prediction methods, including TCRmodel2 and ImmuneBuilder [28], as well as general protein structure prediction methods such as ESMFold and AlphaFold-Multimer [29].

On the most challenging CDR3*β* region, ImmuneFold achieves an RMSD of 1.31 Å (Figure 2**a**, Table S1, Figure S2), outperforming all other methods: TCRmodel2 (1.44 Å), AlphaFold-Multimer (1.57 Å), ImmuneBuilder (1.77 Å), and ESMFold (3.67 Å). Additionally, ImmuneFold attains an RMSD of 1.12 Å on the other CDRs and 0.71 Å on the conserved framework regions, again surpassing the performance of all other approaches.

**Fig. 2:**
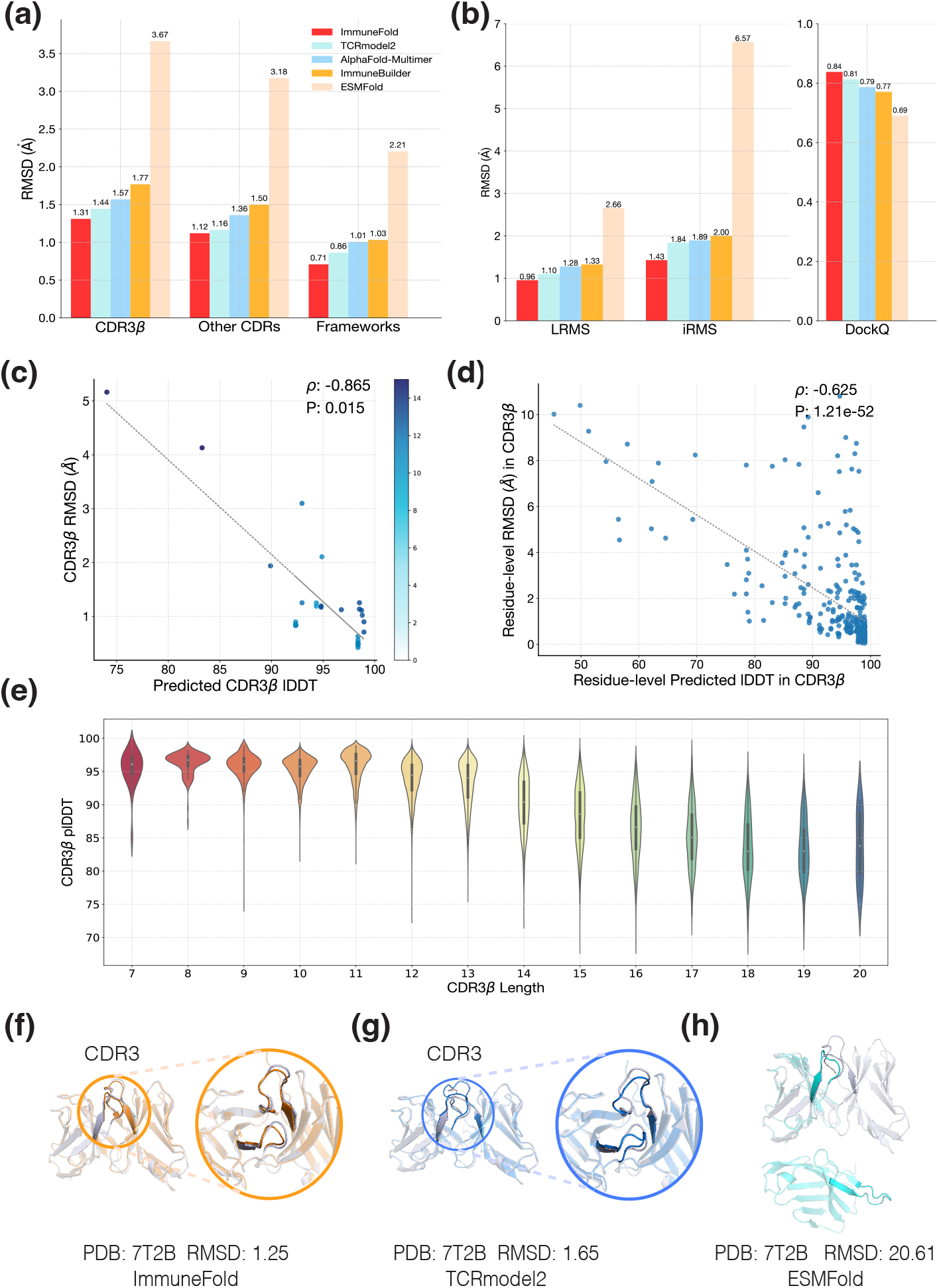
Evaluation of ImmuneFold for TCR structure prediction. **a**, RMSD values for CDR3*β*, other CDRs, and frameworks of predicted TCR structures. **b**, LRMS, iRMS, and DockQ scores of predicted TCR structures. **c**,**d**, Pearson correlation between true RMSD values and predicted lDDT scores of CDR3*β* at the per-target and per-residue level, respectively. **e**, Distribution of predicted lDDT scores for CDR3*β* across various lengths, based on approximately 32,000 predicted TCR structures from the VDJdb dataset. **f-h**, Illustrative examples of predicted TCR structures from ImmuneFold (orange), TCRmodel2 (blue), ESMFold (cyan), and the crystal structure (grey).

We further evaluate the relative poses of the *α* and *β* chains using docking metrics [30] (Figure 2**b**). ImmuneFold achieves an iRMS of 1.43 Å an LRMS of 0.96 Å, and a DockQ score of 0.84, outperforming both TCRmodel2 and AlphaFold-Multimer across all these metrics.

Following AlphaFold2 [6, 31], ImmuneFold also predicts the local distance difference test (pLDDT) between the predicted and ground truth structure as a confidence score. The pLDDT demonstrates high Pearson correlation coefficients of 0.87 and 0.63 with RMSD at the protein and residue levels, respectively (Figure 2**f-g**), indicating that it is a reliable metric for assessing the quality of predicted structures.

We then use `7T2B` as an example to demonstrate the success of our approach (Figure 2**c-e**). In this case, ImmuneFold achieves an RMSD of just 1.25 Å for the entire structure, with no noticeable deviations in the CDRs between the predicted and native structures. In comparison, TCRmodel2 results in a higher RMSD of 1.65 Å and shows clear distortions in the CDRs. ESMFold even fails to determine the correct relative orientation between the *α* and *β* chains, resulting in a significantly larger RMSD of 20.61 Å.

Furthermore, we applied ImmuneFold to predict structures for all the 32,703 TCRs gathered in VDJdb [32] and archived the predicted TCR structures into a database. As of June 2024, only 937 unique TCR-peptide structures have been experimentally determined. Our predicted database represents a 30-fold expansion of the TCR-peptide structure space (Figure 2**i**).

### 2.3 Zero-shot TCR-epitope binding prediction with ImmuneFold

Accurate prediction of TCR-epitope binding is critical for developing immunotherapies, such as neoepitope identification and TCR sequence engineering [1]. We integrate ImmuneFold with the Rosetta software [24] for TCR-epitope binding prediction (Figure 3**a**). It’s important to note that our protocol is a zero-shot learning approach that avoids explicit training on TCR-epitope binding data. In contrast, most existing methods rely on supervised training with limited TCR-epitope binding datasets, leading to poor performance on epitopes not present in the training set [1, 33].

**Fig. 3:**
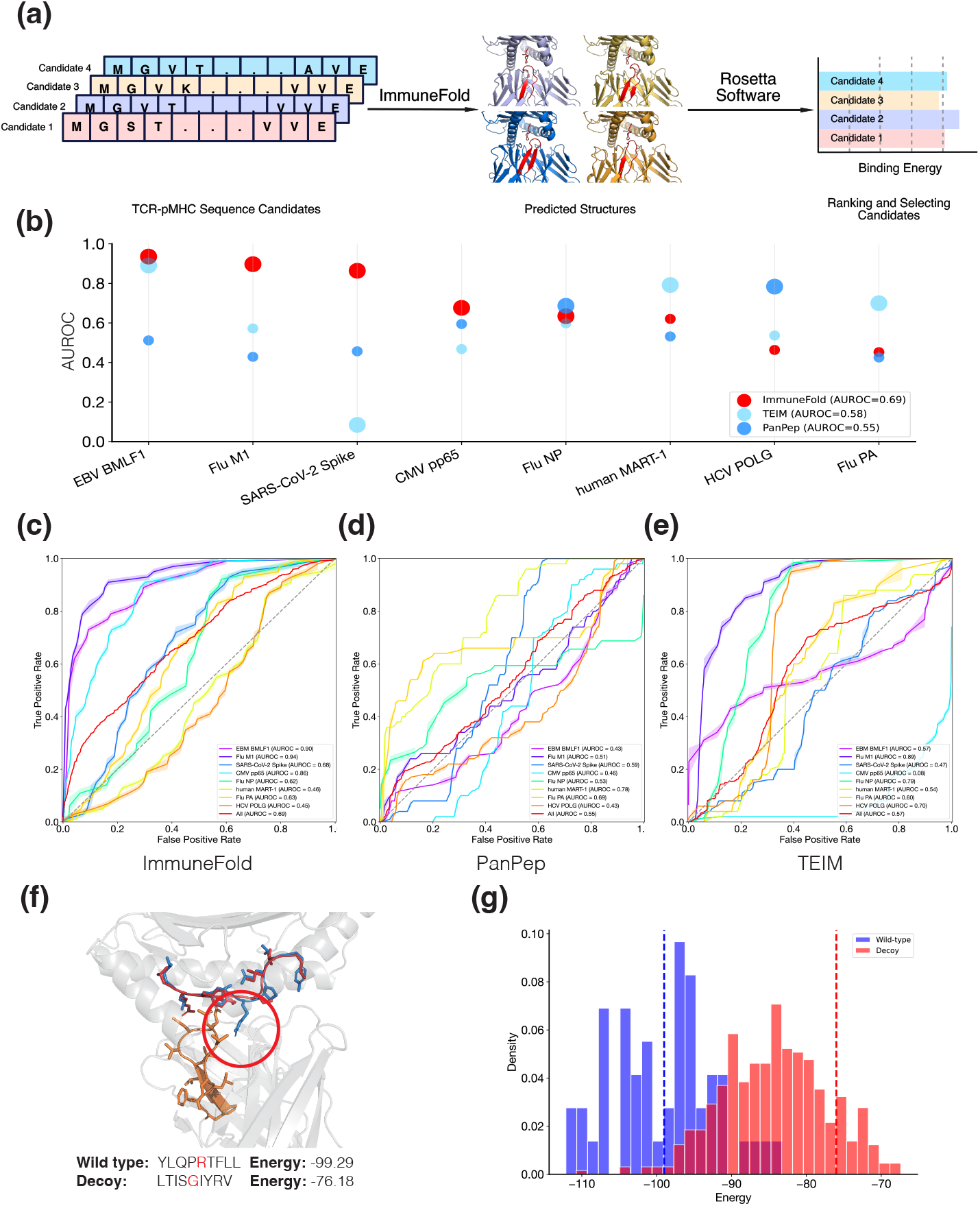
Evaluation of ImmuneFold for TCR-epitope binding prediction. **a**, Framework of TCR-epitope binding prediction with ImmuneFold. **b**, Evaluation results across eight testing subsets, each associated with a wild-type peptide and multiple TCR sequences. Large dots indicate AUROC values that are statistically significant (p-value *<* 0.001, DeLong’s test), whereas small dots represent non-significant results. **c-e**, Receiver operating characteristic (ROC) curves for ImmuneFold, PanPep, and TEIM across the eight testing subsets. **f**, Two examples of TCR-epitope complexes from EBV BMLF1 dataset with significantly different binding energies. The wild-type structure is shown in blue, and the decoy structure in red. **g**, Distribution of Rosetta binding energies for wild-type and decoy structures within the EBV BMLF1 dataset. The blue and red dashed lines represent the energies of the wild-type and decoy complexes in panel **f**, respectively.

To evaluate ImmuneFold, we compiled a TCR-epitope binding dataset comprising eight subsets, each associated with a distinct wild-type peptide and multiple TCR sequences. Wild-type peptides were utilized as positive samples, whereas decoy peptides filtered using NetMHCpan [34] served as negative samples. We then compared the performance of ImmuneFold with that of recent representative TCR-epitope binding prediction methods, specifically PanPep [21] and TEIM [22].

Across these subsets, ImmuneFold achieved the highest AUROC of 0.69 (Figure 3**b-e**), whereas PanPep and TEIM scored 0.55 and 0.57, respectively. Notably, ImmuneFold achieved an AUROC greater than 0.6 with statistical significance (p-value *<* 0.001, DeLong’s test) in 5 out of the 8 testing subsets, while both PanPep and TEIM reached this level of performance in only 2 subsets each.

We then use the EBV BMLF1 subset as an example to investigate the binding energy of TCRs with wild-type and decoy peptides (Figure 3**f**). The interface between the TCR and the wild-type peptide is buried with more interacting residues, whereas the decoy peptide has a mutation to Glycine, resulting in the interface being more exposed to solvent. Consistently, the wild-type exhibits significantly lower Rosetta binding energy than the decoy (Figure 3**g**).

### 2.4 Accurate prediction of antibody structures using ImmuneFold

To assess the performance of ImmuneFold in predicting antibody and nanobody structures, we compiled a test set consisting of 325 antibody and 112 nanobody structures, with no overlap with the training data. We compared ImmuneFold with general protein structure prediction models, ESM-Fold, AlphaFold-Multimer and AlphaFold2, as well as other antibody-specific methods, including IgFold, BALMFold [35], ImmuneBuilder, and tFold-Ab [9].

ImmuneFold is the most accurate in predicting the highly variable CDR H3 region of antibodies, achieving an RMSD of 2.65 Å (Figure 4**a**,**c-d**). The second-best method, tFold-Ab, has an RMSD of 2.99 Å, which is approximately 13% less accurate than ImmuneFold. ImmuneFold also delivers the best performance across the remaining CDRs, the framework regions of antibodies, and almost all regions of nanobodies (Figure 4**a-d**, Figure S3). It is noteworthy that tFold-Ab, IgFold, and BALMFold all rely on language models specifically trained on large antibody sequence datasets, whereas ImmuneFold employs ESM2, a general language model trained on diverse protein families. This highlights the ability of general models to efficiently capture evolutionary constraints in antibodies, consistent with previous studies where antibodies are more successfully designed and validated in wet-lab experiments using general models rather than antibody-specific ones [36, 37].

**Fig. 4:**
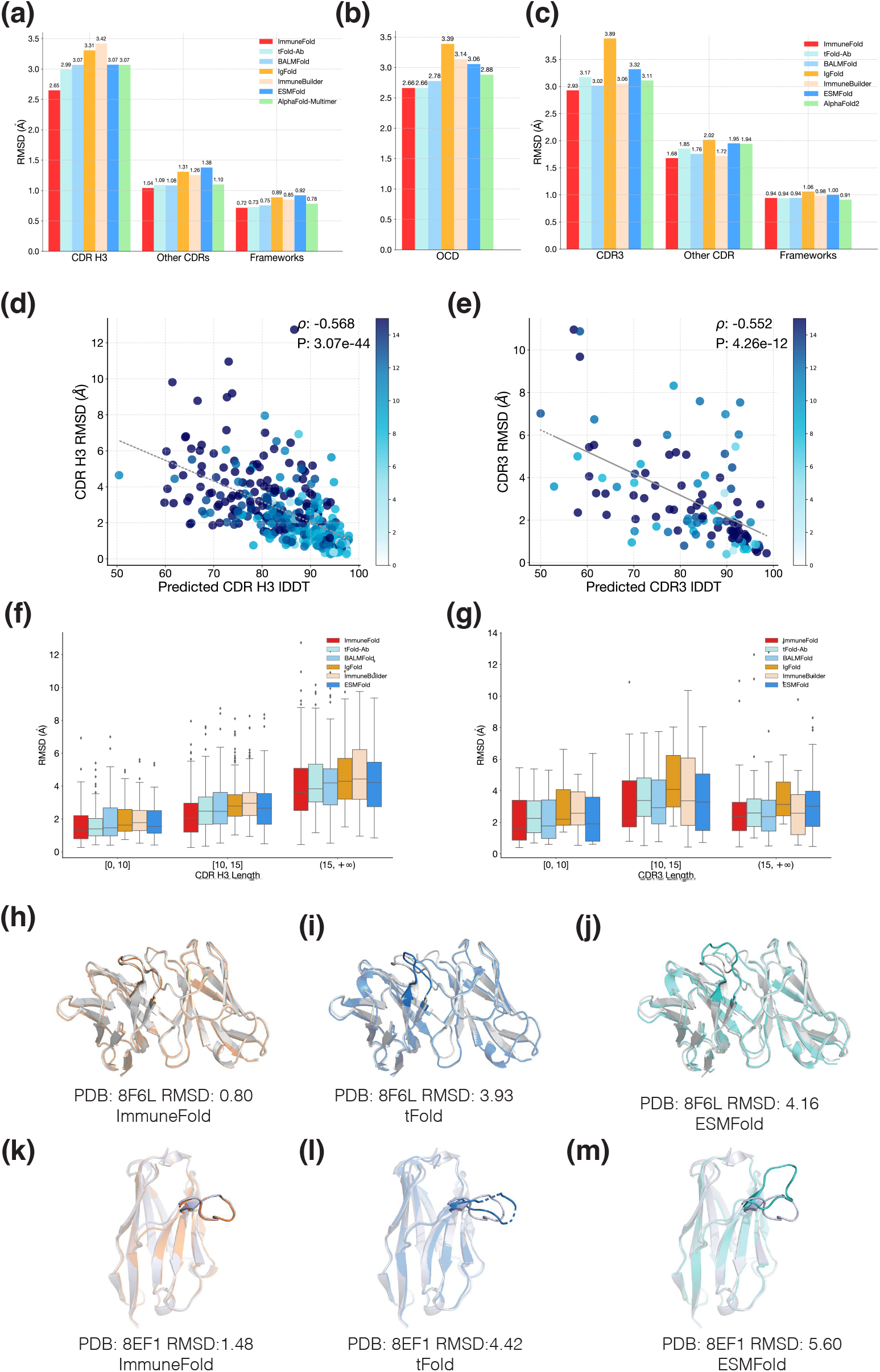
Evaluation of ImmuneFold for antibody and nanobody structure prediction. **a-b**, RMSD and OCD values of predicted antibody structures. **c**, RMSD values of predicted nanobody structures. **d-e**, Pearson correlations between true RMSD values and predicted lDDT of CDR H3 for predicted antibody and nanobody structures, respectively. **f-g**, RMSD values across various lengths of CDR H3 of predicted antibody and nanobody structures. **h-m**, Examples of predicted antibody structures (**h-j**) and nanobody structures (**k-m**) from ImmuneFold (orange), tFold (blue), ESMFold (cyan), and the crystal structure (grey), where the RMSD values on CDR H3 of antibody and CDR3 of nanobody are labeled.

We further evaluate the predicted orientation between heavy and light chains, an important determinant of the overall binding surface. We utilize the orientational coordinates distance (OCD) metric, which summarizes deviations in inter-chain packing angle, inter-domain angle, and heavy-opening and light-opening angles [38]. Among these methods, ImmuneFold and tFold-Ab achieve the most accurate orientation predictions, each with an OCD of 2.66 (Figure 4**a**).

Consistent with the results for TCR structure prediction, ImmuneFold’s predicted lDDT scores exhibit strong Pearson correlations with the true RMSD values at the target level, with coefficients of 0.57 for antibodies and 0.55 for nanobodies (Figure 4**d–e**). Additionally, ImmuneFold achieves consistently superior accuracy across all CDR H3 lengths (Figure 4**f–g**).

We then present two successful examples from ImmuneFold (Figure 4**h-m**). In these cases, the CDR H3 region in the antibody and CDR3 region in the nanobody predicted by ESMFold and tFold-Ab deviate significantly from the native structures and even exhibit steric clashes. In contrast, ImmuneFold provides much more accurate conformations, achieving RMSD values of 0.80 for the antibody CDR H3 and 1.48 for the nanobody CDR3.

### 2.5 Evaluation of antibody and nanobody structure prediction in the presence of antigens

We further investigated how structure prediction accuracy is affected by antigen context. To enable ImmuneFold to consider antigen context during prediction, we provided it with the distogram of the antigen structure as initial recycling features (Methods). We then compared the prediction accuracy of bound antibody and nanobody structures against unbound ones.

When the antigen structure was provided, ImmuneFold reduced the RMSD of predicted antibody CDR H3 from 2.91 Å to 2.57 Å, and that of nanobody CDR3, from 3.00 Å to 2.46 Å (Figure 5**a**,**b**). This indicates its potential application in both antibody virtual screening in the presence of antigen [39, 40] and antibody thermostability optimization in the absence of antigen [41].

**Fig. 5:**
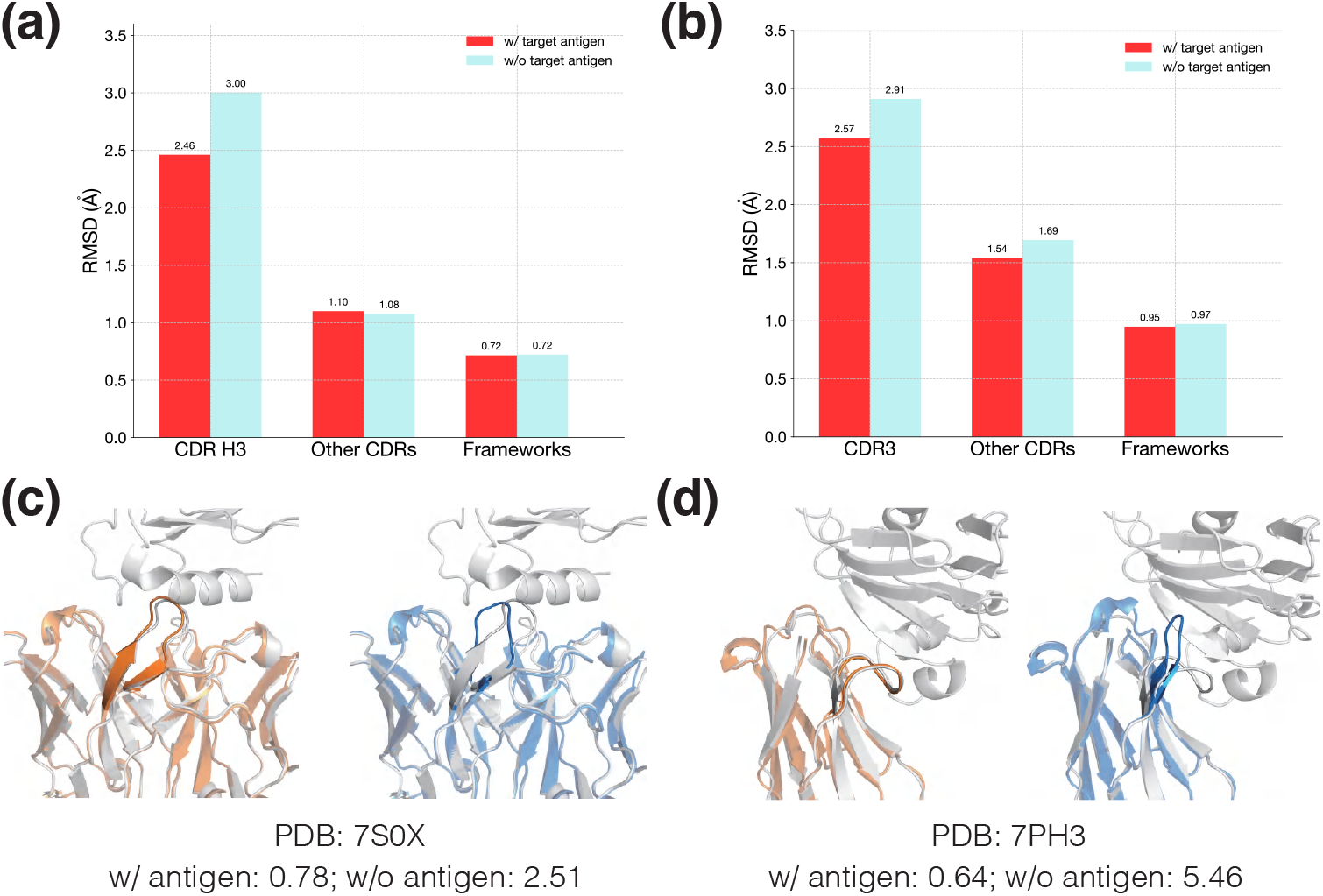
Evaluation of antibody and nanobody structure prediction in the presence of target antigens. **a**,**b**, RMSD values of predicted antibody and nanobody structures when modeled with their target antigens. **c**,**d**, Examples of predicted antibody and nanobody structures, with RMSD values of CDR H3 and CDR3 labeled, respectively. Predictions with antigens are shown in orange, without antigens in blue, and crystal structures are in grey.

We next present two notable cases (Figure 5**c**,**d**) where ImmuneFold significantly improved prediction accuracy with the antigen structure provided,achieving RMSD values of less than 1 Å for the CDR H3 of the antibody and CDR3 of the nanobody.

### 2.6 Training and inference efficiency of ImmuneFold

Previous studies have shown that the parameter-efficient fine-tuning technique LoRA can significantly improve the training efficiency of large-scale Transformer-based models while maintaining prediction accuracy [13, 15]. Here, we investigate the training efficiency of LoRA for fine-tuning Evoformer-like model architectures compared to full-parameter fine-tuning.

We found that LoRA reduces memory usage by nearly half while maintaining almost the same prediction accuracy (Supplementary Figure S1). With the same computational resources and data throughput, LoRA significantly improves training efficiency. This makes it possible for academic labs and other research groups lacking substantial computational resources to fine-tune recently developed, much larger protein structure prediction models [42].

For inference, ImmuneFold has no additional computational overhead compared to ESMFold, as the additional learned parameters are merged into the original parameter matrices after training (Methods). They both take approximately 3 seconds on average to predict a single structure of immune proteins on a Nvidia A100 Card. In comparison, MSA-based methods such as TCRmodel2 [43] and AlphaFold-Multimer [29] usually take approximately 5-20 minutes.

## Discussion

We propose ImmuneFold, a novel method for immune protein structure prediction that efficiently fine-tunes ESMFold specifically on immune protein structures. Our results demonstrate that ImmuneFold outperforms existing methods in predicting the structures of various immune proteins, including T-cell receptors, antibodies, and nanobodies. Notably, when integrated with Rosetta software, ImmuneFold can accurately predict TCR-epitope binding, indicating its potential for real-world applications in immunotherapy.

ImmuneFold and other recent deep learning–based methods [8, 28, 35] only predict static structures of immune proteins. However, T-cell receptors and antibodies can exhibit conformational changes upon binding to specific antigens [44–50]. ImmuneFold can be extended in two ways to address this challenge. First, incorporating molecular dynamic simulation data for immune proteins during training could enable the model to better capture the conformational diversity of these proteins. Second, our strategy can be extended to fine-tune generative models for protein structure prediction, allowing protein conformations sampling rather than predicting a single static structure [42, 51].

While we have demonstrated that ImmuneFold improves TCR–epitope binding prediction, further efforts are needed to enhance antibody-antigen docking performance, especially when the antigen structure is large and the epitope on the antigen is unknown. Simply fine-tuning ESMFold is insufficient for this task. Antibodies can target a wide range of foreign proteins and may form considerably large antibody-antigen complexes, whereas TCRs typically target peptides of only 10–20 amino acids. However, ESMFold and its underlying language model, ESM-2, are trained only on monomeric proteins, making it difficult to capture the inter-chain evolutionary constraints critical for modeling large protein complexes [12, 52]. Our future work will primarily focus on improving antibody-antigen docking performance by combining the advantages of ImmuneFold with other cutting-edge methods. For instance, we will explore integrating ImmuneFold with physical energy-based docking methods, as adopted in previous studies [9, 53], or enrich our training dataset with docked protein complex structures [40, 54].

## 3 Methods

### 3.1 Datasets

#### TCR structure datasets

We derived the training and test sets for TCR structure prediction from the STCRDab database [27]. We excluded MHC Class II complex structures, antigen chains whose antigen type was not a peptide, and TCRs lacking the *β* chain. The dataset was partitioned into training and test sets based on the release date of October 1, 2022. Following previous work [43], we remove the overlap between the training and testing set using the threshold of 99% *β* sequence similarity for a fair comparison. As a result, the final training and test sets comprised 730 and 27 complexes, respectively.

#### Antibody and nanobody structure datasets

For antibody and nanobody structure prediction, we construct training and test sets from the SAbDab database [11]. The dataset is split into training and testing sets based on the release date of 1 January 2022. We remove the overlap between the training and testing set using a threshold of 90% sequence similarity. Additionally, we applied a 90% sequence identity threshold within the test set to remove redundant sequences. The final dataset includes a total of 9829 antibody and nanobody structures for training, and 325 antibodies and 112 nanobodies for testing, respectively.

#### TCR-epitope binding dataset

Following the strategy of previous methods [21, 55], we compiled a testing set for TCR-epitope binding prediction, consisting of 382 positive and 3,438 negative samples. Specifically, the testing set includes eight subsets, each associated with a distinct wild-type peptide and multiple TCR sequences. These wild-type TCR-peptide complexes were utilized as positive samples. For each positive sample, we filtered and selected nine decoy peptides of equivalent lengths using NetMHCpan [34].

#### Large-scale TCR sequence data for structure prediction

We collected a dataset of 32,703 TCR-peptide sequences from the VDJdb database [32], including only sequences associated with MHC Class I complexes.

### 3.2 Parameter-efficient fine-tuning on ESMFold

ESMFold [12] leverages a large-scale protein language model for protein structure prediction and consists of three main components: a large protein language model, the `Folding Trunk module`, and the `Structure module`. The Folding Trunk is an Evoformer-like architecture adapted from AlphaFold, designed to learn single and pairwise representations of evolutionary and geometric constraints.

In our approach, we fine-tune ESMFold on immune protein structures using Low-Rank Adaptation (LoRA) [13]. Specifically, we apply LoRA to the weight matrices of the `Linear` layers within the Folding Trunk, including the `AttentionWithPairBias, OutProductMean, Transition`, and `TriangularUpdate` modules [6, 12]. All parameters of the language model and the structure module are kept frozen during this process.

LoRA restricts parameter updates to a low-rank matrix Δ*W*, which is added to the original weight matrix *W*_0_ ∈ ℝ^*m×n*^. This low-rank matrix is factorized into two smaller matrices *B* ∈ ℝ^*m×r*^ and *A* ∈ ℝ^*r×n*^, where *r ≪ m* and *r ≪ n*. The forward computation is expressed as:

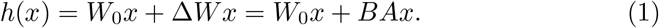

Here, *B* is initialized as zeros, and *A* is initialized from a `Kaiming Uniform` distribution [56]. After training, the learned *A* and *B* are merged into *W*_0_, resulting in no additional computational overhead for inference.

We set the rank *r* to 16 for single representations and 8 for pair representations. This configuration reduces the number of trainable parameters to 2.7% of that required for full-parameter fine-tuning, enabling efficient adaptation of ESMFold to immune protein structures with minimal computational cost.

Following ESMFold, we employ an offset of 512 for relative positional encoding between different chains, supplying these as features to the `Evoformer` module. However, unlike ESMFold, which adds a linker of 25 glycines before extracting ESM2 language embeddings, we adapt the code of the original ESM2 model to handle position offsets directly.

### 3.3 Immune protein structure prediction with antigen context

ImmuneFold supports structure prediction of bound antibodies or TCRs in the presence of specific antigens. We adapt the original ESMFold architecture to incorporate antigen structure during the network recycling stage.

AlphaFold2 [6] and subsequent methods like ESMFold [12] utilize network recycling to iteratively refine predicted structures, significantly improving prediction accuracy. In this process, the distogram of residue-residue distances and other features from the previous predictions are fed back into the model, providing feedback for further refinement.

For ImmuneFold, in the first recycling cycle, we initialize the antibody or TCR distogram with zeros and compute the distogram for the antigen structure directly. In subsequent cycles, the antibody or TCR distogram is updated with the latest predictions, while the antigen’s distogram remains unchanged.

### 3.4 Training losses and other hyper-settings

Following AlphaFold-Multimer [29], we incorporate the interface Frame Aligned Point Error (FAPE) into the original loss functions used by ESMFold. The training loss is expressed as:

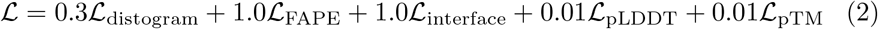

We train a single model for antibody and nanobody structure predictions. To adapt to various application scenarios, we randomly remove either the light chain or the antigen chain in antibody-antigen complex structures during training. To accommodate cases where the antigen structure is available in the application, we replace the predicted antigen structure with the ground truth during the calculation of distogram features in the recycling stage [6].

For training in the TCR dataset, we augment the data by randomly removing the peptide chain, the MHC chain, or both.

We utilize the Adam optimizer [57] with an initial learning rate of 5 *×* 10^*−*4^. Each model is trained for 2*×*10^4^ steps with a batch size of 32 structures. LoRA fine-tuning requires *∼* 10 days per model using eight Nvidia A800 GPUs, whereas the full-parameter fine-tuning under the same training settings requires *∼*20 days per model.

### 3.5 TCR-epitope binding prediction

For TCR-epitope binding prediction, we first predict the complex structure using ImmuneFold and then evaluate the binding energies using Rosetta software. We directly compute the binding free energy (Δ*G*) of the predicted complex structures, following a method similar to that used in ATLAS [58]:

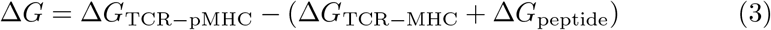

where Δ*G*_TCR*−*pMHC_ denotes the free energy of the entire TCR-pMHC complex, computed using the `ref2015` scoring function [58] in PyRosetta [59]. The terms Δ*G*_TCR*−*MHC_ and Δ*G*_peptide_ represent the energies for isolated structures of peptide and TCR-MHC, respectively.

### 3.6 Evaluation metrics

To evaluate the performance of immune structure predictions, we compute the root mean squared deviation (RMSD) between predicted and experimentally determined structures, focusing solely on backbone heavy atoms (N, C_*α*_, C, O). Specifically, we align the entire variable (Fv) regions of the predicted and crystal structures using the `Kabsch algorithm` [60], then compute the RMSD separately for each CDR and framework region. We define the CDR and framework regions using the IMGT numbering scheme [61] for TCR structures and the Chothia numbering scheme [62] for antibody and nanobody structures.

To assess the relative positioning between the *α* and *β* chains in TCRs, we employ the interface RMSD (iRMS), ligand RMSD (LRMS), and DockQ metrics [30, 63]. For antibodies, we utilize the Orientational Coordinate Distance (OCD) to evaluate the relative position between heavy and light chains [38].

For the evaluation of TCR-epitope binding prediction, we use AUROC to assess sorting accuracy. AUROC measures the area under the curve with the false positive rate (FPR) on the x-axis and the true positive rate on the y-axis.

## Data Availability

The training and test datasets utilized in this study are derived from the STCRDab (https://opig.stats.ox.ac.uk/webapps/stcrdab-stcrpred/) and SAbDab (https://opig.stats.ox.ac.uk/webapps/sabdab-sabpred/sabdab) databases. Data of TCR-epitope binding prediction is obtained from the link (https://github.com/phbradley/TCRdock/tree/main/datasets from the paper). Additionally, the VDJdb database is accessed via its website (https://vdjdb.cdr3.net/).

## Code Availability

The ImmuneFold software is available on GitHub at https://github.com/CarbonMatrixLab/immunefold.

## Acknowledgments

We acknowledge the financial support from the National Natural Science Foundation of China (grant no. 32370657) and the Project of Youth Innovation Promotion Association CAS to H.Z. We also acknowledge the financial support from the Development Program of China (grant no. 2020YFA0907000) and the National Natural Science Foundation of China (grant nos. 32271297 and 62072435) to D.B. We thank Beijing Paratera Co., Ltd and the ICT Computing-X Center for providing computational resources.

We also thank Dr. Yufeng Shen and Chungong Yu for the useful discussions on our work.

## Author Contributions

H.Z. conceived the project and reimplemented ESMFold with LoRA integration. T.Z. and M.R. adapted the training procedures for structure prediction of antibody-antigen complexes and TCR–pMHC complexes, and developed the zero-shot protocol for TCR–epitope binding prediction. T.Z., M.R., and Z.H. conducted the experiments and wrote the manuscript. H.Z., D.B., S.T., and M.L. revised the manuscript.

## 4 Supplementary Materials

### 4.1 Details on running the compared methods

*ImmuneBuilder*: We utilized the test script provided in the ImmuneBuilder GitHub repository (https://github.com/oxpig/ImmuneBuilder) and conducted TCRBuilder2, AbBuilder2, and NanoBuilder2 for TCR, antibody, and nanobody structure prediction, respectively, using the default settings.

*TCRmodel2*: We used the test script provided in the TCRmodel2 GitHub repository (https://github.com/piercelab/tcrmodel2). For a fair comparison, we employed AlphaFold-Multimer in TCRmodel2 with only model 1, rather than an ensemble of 5 models, and set the template time cutoff to 1 October 2022.

*AlphaFold-Multimer*: We utilized the test script provided in the AlphaFold repository (https://github.com/google-deepmind/alphafold). For a fair comparison, we used AlphaFold-Multimer with only model 1 and set the template time cutoff to 1 January 2022 for antibodies and 1 October 2022 for T-cell receptors.

*AlphaFold2*: We used the test script from the AlphaFold repository (https://github.com/google-deepmind/alphafold). For a fair comparison, we utilized only model 1 and set the template time cutoff to January 1, 2022.

*ESMFold*: We utilized the test script provided in the ESM GitHub repository (https://github.com/facebookresearch/esm) with the default settings.

*IgFold*: We utilized the test script provided in the IgFold GitHub repository (https://github.com/Graylab/IgFold) with the default settings.

*BALMFold*: We utilized all predicted structures provided by the BALM-Fold server (https://beamlab-sh.com/models/BALMFold).

*tFold-Ab*: We utilized the test script provided in the tFold GitHub repository (https://github.com/TencentAI4S/tfold) with all default parameters.

*PanPep*: We utilized the test script provided in the Panpep GitHub repository (https://github.com/bm2-lab/PanPep) in the zero-shot setting.

*TEIM*: We utilized the test script provided in the TEIM-seq GitHub repository (https://github.com/pengxingang/TEIM) with the sequence-level interaction.

## Supplementary Figures

**Fig. S1:**
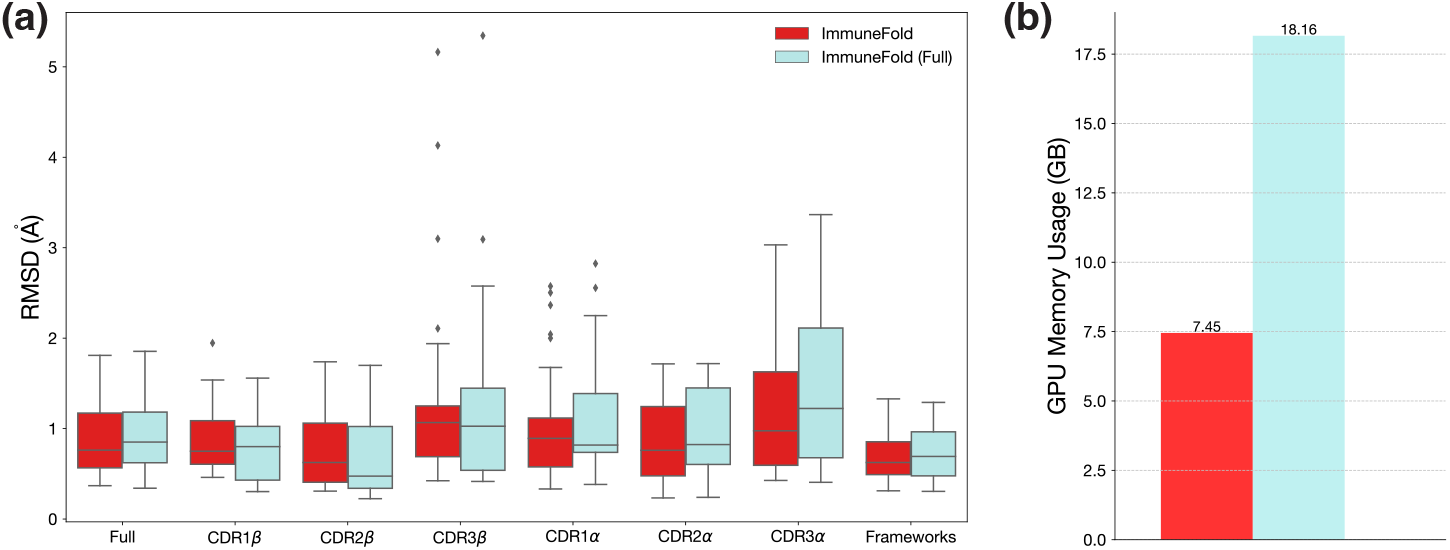
Evaluation of ablation study. **a**, Evaluation on RMSD for CDR3*β*, other CDRs, and frameworks for the TCR test sets. **b**, Evaluation of GPU memory usage under different configurations.

**Fig. S2:**
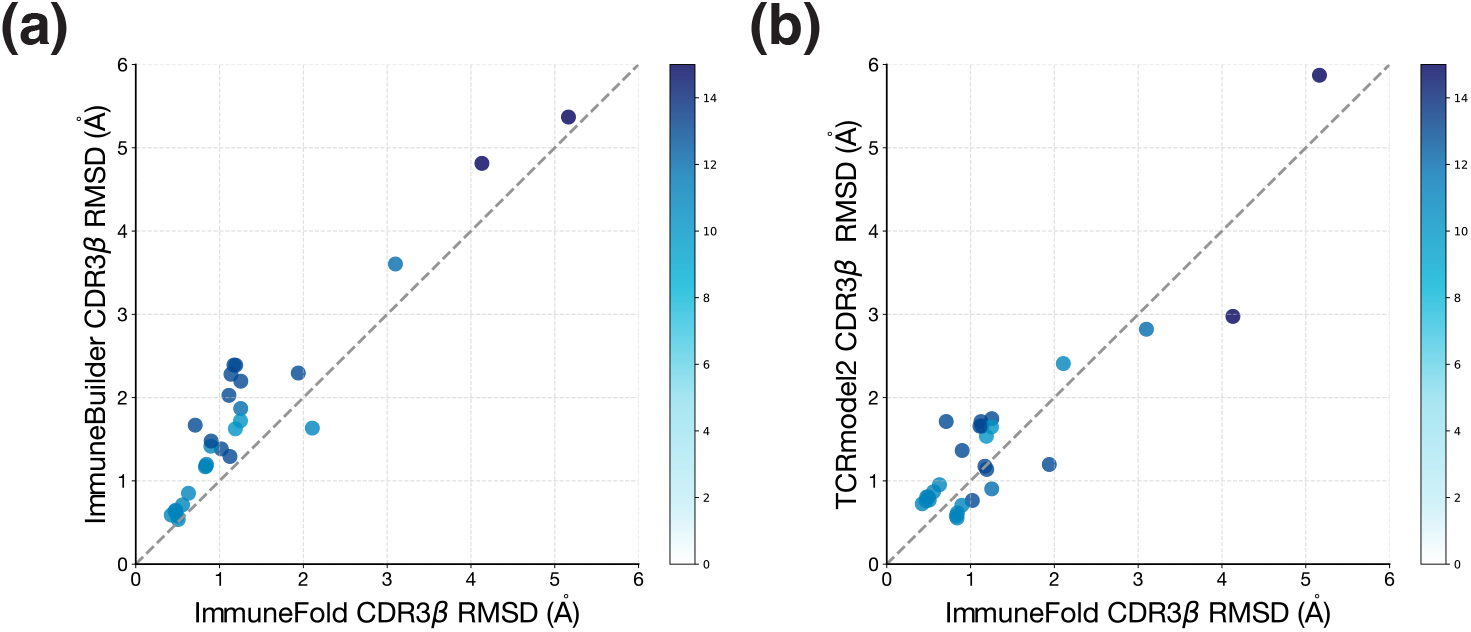
Head-to-head comparison with ImmuneBuilder and TCRmodel2 on T-cell receptor dataset.

## Supplementary Tables

**Table S1:** TCR structure prediction accuracy.

| Methods | CDR1β | CDR2β | CDR3β | CDR1α | CDR2α | CDR3α | Framework |
| --- | --- | --- | --- | --- | --- | --- | --- |
| ESMFold | 2.69 | 3.10 | 3.67 | 2.66 | 2.97 | 3.44 | 2.21 |
| ImmuneBuilder | **0.85** | 0.90 | 1.77 | 1.35 | 1.04 | 1.92 | 1.03 |
| TCRmodel2 | 0.93 | **0.75** | 1.44 | **0.94** | 1.00 | 1.42 | 0.86 |
| AlphaFold-Multimer | 1.05 | 0.90 | 1.57 | 1.13 | 0.98 | 1.70 | 1.01 |
| ImmuneFold | 0.87 | 0.76 | **1.31** | 1.06 | **0.88** | **1.27** | **0.71** |

**Fig. S3:**
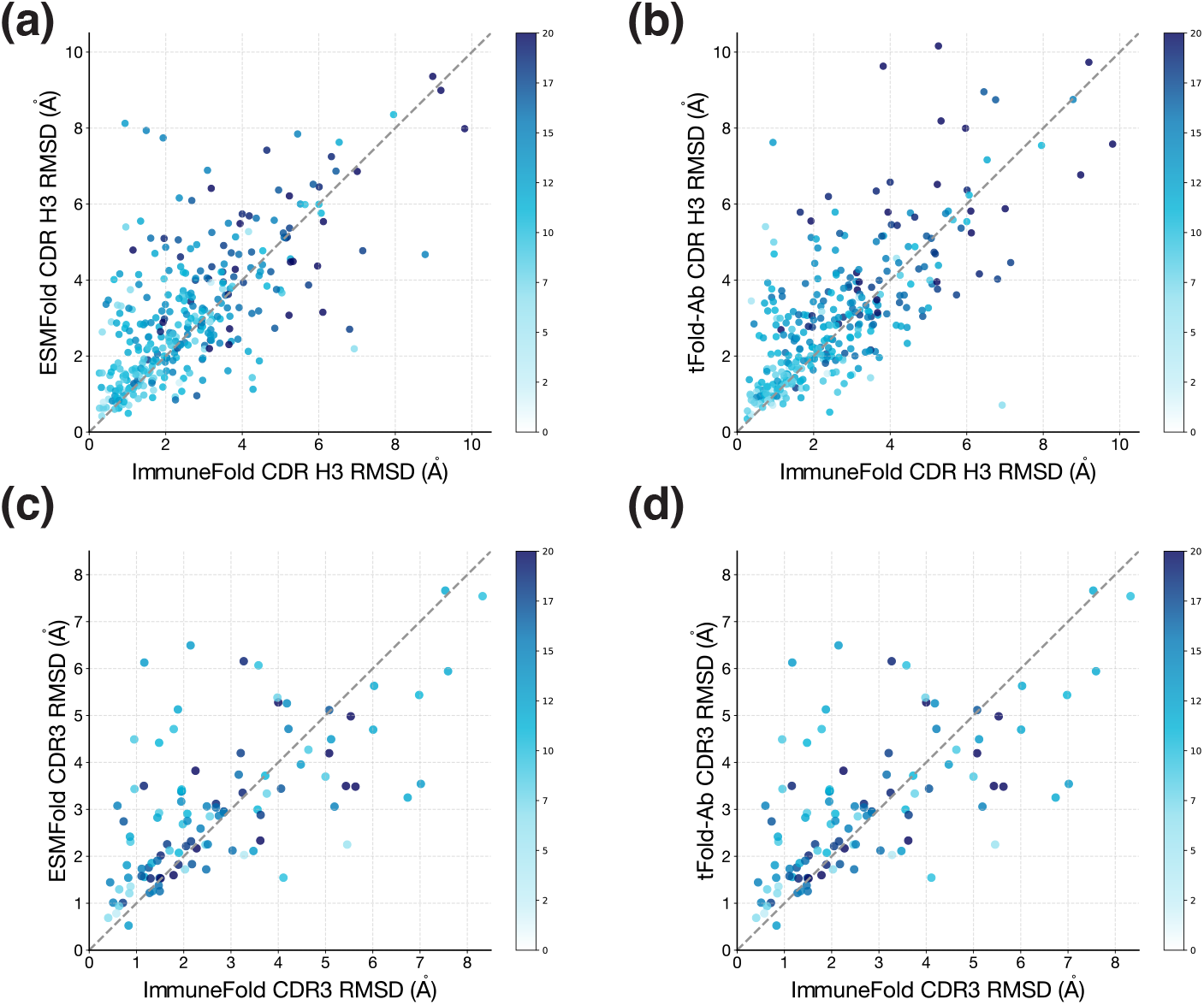
Head-to-head comparison with representative methods. **a-b**, Results on antibody dataset. **c-d**, Results on nanobody dataset.

**Table S2:**
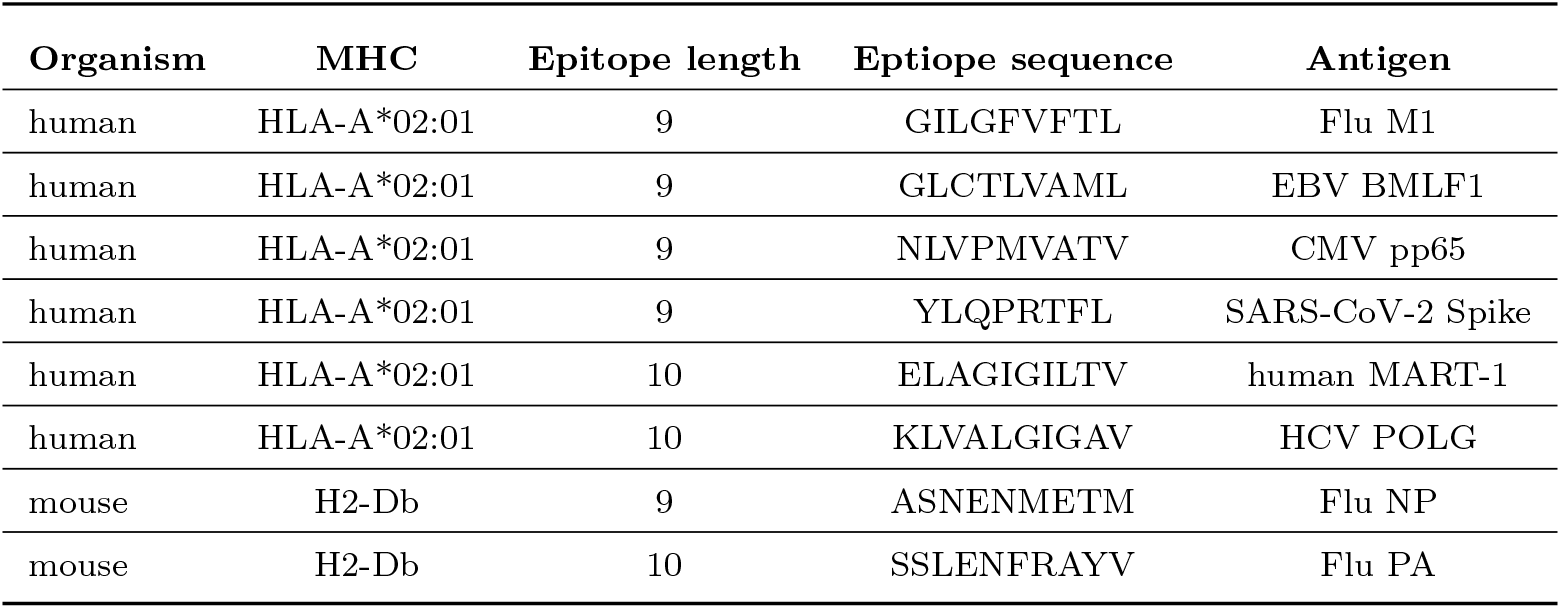
Neoepitope benchmark.

**Table S3:** Antibody structure prediction accuracy.

| Methods | CDR H1 | CDR H2 | CDR H3 | CDR L1 | CDR L2 | CDR L3 | Framework |
| --- | --- | --- | --- | --- | --- | --- | --- |
| ESMFold | 1.37 | 1.33 | 3.07 | 1.24 | 0.88 | 1.36 | 0.92 |
| AlphaFold-Multimer | 1.16 | 1.05 | 3.07 | 0.91 | 0.77 | 1.08 | 0.78 |
| ImmuneBuilder | 1.38 | 1.23 | 3.42 | 1.03 | 0.93 | 1.20 | 0.85 |
| IgFold | 1.38 | 1.22 | 3.31 | 1.10 | 0.91 | 1.39 | 0.89 |
| tFold-Ab | 1.20 | 1.04 | 2.99 | 0.88 | 0.77 | 1.06 | 0.73 |
| BALMFold | 1.18 | 1.03 | 3.07 | 0.86 | 0.76 | 1.08 | 0.75 |
| ImmuneFold | **1.14** | **1.00** | **2.65** | **0.65** | **0.75** | **1.00** | **0.72** |

**Table S4:**
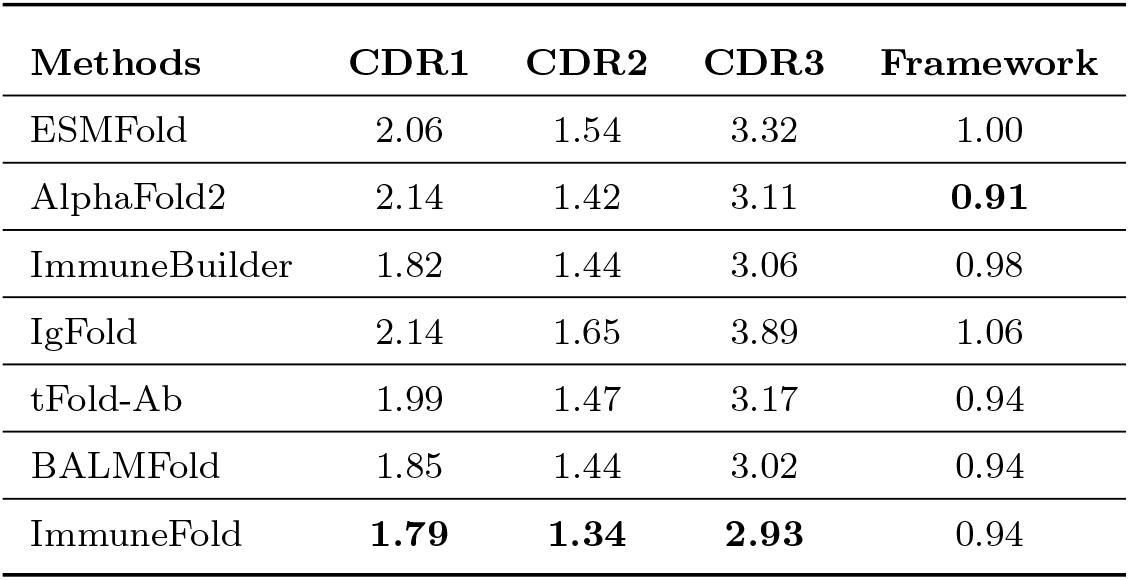
Nanobody structure prediction accuracy.

| Methods | CDR1 | CDR2 | CDR3 | Framework |
| --- | --- | --- | --- | --- |
| ESMFold | 2.06 | 1.54 | 3.32 | 1.00 |
| AlphaFold2 | 2.14 | 1.42 | 3.11 | **0.91** |
| ImmuneBuilder | 1.82 | 1.44 | 3.06 | 0.98 |
| IgFold | 2.14 | 1.65 | 3.89 | 1.06 |
| tFold-Ab | 1.99 | 1.47 | 3.17 | 0.94 |
| BALMFold | 1.85 | 1.44 | 3.02 | 0.94 |
| ImmuneFold | **1.79** | **1.34** | **2.93** | 0.94 |

**Table S5:** Antibody structure prediction accuracy given antigen contexts.

| Methods | CDR H1 | CDR H2 | CDR H3 | CDR L1 | CDR L2 | CDR L3 | Framework |
| --- | --- | --- | --- | --- | --- | --- | --- |
| w/ antigen | 1.11 | 0.98 | 2.46 | 0.95 | 0.71 | 1.03 | 0.72 |
| w/o antigen | 1.18 | 1.08 | 3.00 | 0.86 | 0.76 | 1.05 | 0.72 |

**Table S6:** Nanobody structure prediction accuracy given antigen contexts.

| Methods | CDR1 | CDR2 | CDR3 | Framework |
| --- | --- | --- | --- | --- |
| w/ antigen | 1.65 | 1.25 | 2.57 | 0.95 |
| w/o antigen | 1.81 | 1.34 | 2.91 | 0.97 |

